# *Citius, Fortius?* Cohort, inflammation and trajectories of gait speed and grip strength in older Britons

**DOI:** 10.1101/268334

**Authors:** Gindo Tampubolon, Maria Fajarini

**Affiliations:** Manchester Institute for Collaborative Research on Ageing University of Manchester; Evidence & Analytics, Manchester

**Keywords:** English Longitudinal Study of Ageing, gait speed, grip strength, inflammation, C-reactive protein, fibrinogen, attrition, weighting

## Abstract

Although the cost of long term care of physical disabilities is considerable, little is known about individual trajectories of physical function (measured by gait speed and grip strength) that preceded the process of disablement. Moreover, studies on trajectories of health function have often ignored cohort composition, precluding evidence of secular improvement. And few have explored the role of chronic inflammation on older people’s physical function trajectories. Using the English Longitudinal Study of Ageing 2004–2013 we derived trajectories of gait speed and grip strength of Britons aged ≥ 50 years and investigated the effect of inflammation. Then we drew trajectories for different cohorts to seek evidence of secular improvement. We uncovered a complex gradient of improvement in trajectories of physical function that depends on sex and maximum versus normal capacity. In conclusion, accounting for the cohort composition of older people can materially modify the future cost of long term care.

## Introduction

Olympians are not the only group attaining secular improvement over recent decades. The most recent cohort of older people have also showed higher levels of cognitive function [1]. The Post-War cohort, compared to the earlier cohorts, maintained higher episodic memory at similar ages. Given the myriad connections between cognitive and physical functions [2], this evidence raises a possibility that the more recent cohort of older people also maintain an advantaged trajectory of physical function. We interpret inter-cohort (for instance from War cohort to Post-War cohort) increase in functioning as evidence of secular improvement [3].

We tested this possibility by using gait speed and grip strength as two well-characterised measures of physical function [4–6], especially since grip strength has been shown to predict disability, morbidity, and mortality [7, 8].

Such a possibility can have theoretical and practical implications. Cohort has always been an important concept waiting to take its place in our scheme of understanding individual change over an extended period of later life (≥ 50 years). Cohort in our sense is not primarily defined chronologically. Instead it refers to a time course or an era defined by socio-historical events, e.g. War vs. Post-War cohort or Pre-Depression era vs. Depression era cohort. But it has rarely been treated as an integral part, as if by later life age has chiseled off any cohort difference that might matter.

However, recent scholarship has questioned this age-as-leveller assumption. Birth cohort, as a marker for a childhood stage exposed to similar kinds of developmental hazards and damage (cf. the War cohort versus the Post-War cohort), can give insight into how later life experience might unfold. For example, in the English Longitudinal Study of Ageing (ELSA), childhood condition at age ten has been found to exert a very long influence indeed. Those who were poor financially in their first decade of life displayed slower gait, poorer memory, and more depression in their fifth to ninth decades of life [5]. Because the evidence is based on cross-sectional observations, the study cannot distinguish cohort effect from age trajectory effect. It remains a distinct possibility that a net cohort effect yields insight on how an early stage in life course shapes health outcomes and wellbeing throughout later life.

The implications are not merely theoretical [9]. In practice, cohort effect can have a bearing on policy to support and care for older population, a growing and important demographic. For example, two British regions with the same number of people aged ≥ 50 years but different cohort composition are likely to yield different evolution of demand for health and social care. To put a figure on the stake, the Swedish and Dutch governments spent 3.5% of their gross domestic products on long term care of older people with disabilities, including physical disabilities (WHO World Report on Ageing and Health) [10]. Apart from the difference in their health systems, difference in cohort composition across the two countries can impact future trends of these percentages.

The WHO World Report also emphasised the need for refined pictures of changes in physical function of individuals in later life. This calls for deriving, based on the experience of older people living in communities or outside institutions, age trajectories of physical function over an extended period.

One note of caution follows when deriving such trajectories. Collecting repeated measures from older people inevitably faces attrition problems, since older people tend to attrite non-randomly in subsequent visits [11, 12]. Recently, a number of solutions have been proposed including weighting and joint modelling [1, 13, 14]; inverse proportional to attrition weighting is applied here.

While recent studies on cognitive function and well-being showing cohort improvement have prompted the question on cohort effect [1, 13], another recent study on blood-based biomarkers of cognitive deficits [15] have suggested another insight. The study showed that inflammation, measured using two inflammatory markers of high sensitivity C-reactive protein (CRP) and fibrinogen, is associated with cognitive deficits. So given the connections between cognitive and physical functions [2], inflammation can be expected to have an effect on physical function trajectories throughout later life as well [16]. This can enrich our understanding of key drivers of healthy ageing that affect multiple dimensions of health functions.

We therefore aimed to gauge whether there is a secular improvement to gait speed and grip strength enjoyed by some older people living today and to quantify the effect of inflammation on the trajectories of physical function. We note that our aim was not to estimate causality, merely to explore important association hitherto neglected. To tie the strands together we raise three questions: What is the shape of age trajectories of physical function as measured by gait speed and grip strength? Do recent cohorts possess a more advantaged trajectory, one with higher levels at similar ages to the earlier cohorts? Is inflammation associated with lower levels of physical function?

## Materials and methods

The University of Manchester’s institutional review board has exempted this study since it used publicly available anonymised secondary data for research.

The English Longitudinal Study of Ageing (ELSA) is the main resource for a nationally-representative ageing study of the English older population. The first wave was in 2002 and subsequent waves follow biennially. Repeated biomarker information is available from the even numbered waves (2004/5, 2008/9 and 2012/3) when nurses visited the participants. The data are freely available from the UK Data Archive (www.data-archive.ac.uk) as study number 5050. More details of the study are given elsewhere [14, 17–19].

We used two measures of physical function that have been used in this sample [4, 5]. The first is gait speed at normal pace (m/s), timed by a research nurse, in m/s; the second is objectively measured maximum grip strength of the dominant hand (kg), obtained using a dynamometer (Smedley Dynamometer, Tokyo). The nurse demonstrated each test before the participant was asked to do it. After adjusting the dynamometer to suit the participant’s hand and positioning the participant correctly, the participant was asked to squeeze the dynamometer as hard as possible for a couple of seconds. Three values were recorded for each hand, starting with the non-dominant hand and alternating between hands. The maximum was used. Walking speed was measured by marking a course in a suitable space in the participant’s home with a tape measure and placing masking tape at the starting and ending points. The length of the course was 244 cm. The nurse used a stopwatch to record the time. The participant was asked to complete two timed walks at normal pace.

Blood samples were collected by the research nurse in three waves and kept deep-frozen until analysis at the Newcastle NHS hospital laboratory. Plasma samples were analyzed for fibrinogen concentrations (in g/L) using an ACL TOP CTS analyzer. The samples were also analyzed for high sensitivity CRP concentrations (in mg/L), applying a particle-enhanced immunoturbidimetric assay, using Roche Modular P analyzer. Both have been used before [15]. Following the literature [15, 20, 21], we removed observations with CRP concentrations above 10 mg/L, indicating acute inflammation. Because CRP distribution is skewed, following the Women’s Health Initiative Study and Established Populations for Epidemiologic Studies of the Elderly, we derived an indicator variable marking concentrations in the top quartile; similarly with fibrinogen [22–24]. We constructed four cohorts or birth groups marked by socio-historical events, to make them comparable to the US sister study (Health and Retirement Survey) and a previous study of this sample [1]. The four cohorts (requiring three cohort indicators) are pre-Depression cohort (born before 1930, omitted as the reference), Depression era cohort (1931–1938), War cohort (1939–1945) and post-War cohort (born after 1946). These four are more refined than the three cohorts used in the ELSA report [17, 25, 26].

Following the literature [4, 27] we built separate models for the sexes. We used mixed model to derive trajectories of both physical function measures; the model is variously known as latent growth curve model or random coefficients model. The dependent variables were gait speed and grip strength. The key covariates were cohort indicators, high CRP indicator, and high fibrinogen indicator. In applying mixed or random coefficients model, we estimated random intercepts, random slopes of age, and their covariance. As confounders we included, in addition to age and sex other covariates including wealth (in tertiles with the poorest third as reference), marital status (single, married or partnership, and others as reference) and socioeconomic positions earlier in the life course: education (threefold: less than high school as reference, high school, and college), and occupational class (threefold: routine manual as reference, intermediate, and managerial/professional). We also included heigh following similar studies in Europe and Britain [28–31]. We set an a priori *α* = 0.05.

We retained in the analysis those with complete information on both
physical function measures (gait speed and grip strength), both inflammatory markers (CRP and fibrinogen) and other covariates (as analytic sample). This resulted in *N* = 5,030 at baseline, and 5,384 and 4,500 at subsequent waves. The analytic and reference samples were tested for difference using *t* test (continuous covariates) and *χ*^2^ test (nominal covariates). At baseline, the analytic sample (compared to the reference sample) had relatively younger participants (66.0 [standard deviation/SD 9.1] vs 69.6 [SD 9.9]), *t* = 12.4, *p* < 0.001, lower CRP levels (2.6 [SD 2.3] vs 2.9 [SD 2.3]), *t* = 2.3, *p* = 0.020), and lower fibrinogen levels (3.1 [SD 0.6] vs 3.3 [SD 0.9], *t* = 3.5, *p* < 0.001), higher gait speed (1.1 [SD 0.3] vs 1.0 [SD 0.4] m/s, *t* = 5.4, *p* < 0.001), and stronger [SD 10.7] vs 28.2 [SD 10.8] kg, *t* = 2.4, *p* = 0.015). higher proportion of women to men (analytic sample: 0.53, SD 0.49; reference sample: 0.56, SD 0.49) but there was no significant difference between the samples 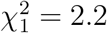, *p* = 0.14.

Because the repeated observations had shrunk due to attrition, we followed the extensive literature in using inverse proportional to attrition weighting [32–35]. Particularly following [35], the attrition model included age, sex, smoking, cognition, education, hypertension, cardiovascular disease, diabetes, and retirement status; then stabilised weights were computed with base model including age, sex, and education. All modelling was done in Latent GOLD Syntax 5.1 [36].

## Results

Women made up the majority of the sample (8,142; 54.6%) while the average gait speed at normal pace was 1.1 m/s (standard deviation [SD] 0.4 m/s), of maximum grip strength was 29.5 kg (SD 11.5 kg), and of age was 66.4 year (SD 8.9 yr); see Table 1.

**Table 1.** Description of analytic sample with reference categories in asterisks. Source: ELSA 2004-2013.

| Variable | mean or $N$ | Std. Dev. or % |
| --- | --- | --- |
| Gait speed | 1.1 | 0.4 |
| Grip strength | 29.5 | 11.5 |
| CRP | 2.5 | 2.3 |
| Fibrinogen | 3.1 | 0.6 |
| Age, year | 66.4 | 8.9 |
| Men | 6,772 | 45.4 |
| Women | 8,142 | 54.6 |
| Marital status |  |  |
| Single | 830 | 5.6 |
| Married | 10,272 | 68.9 |
| Separated/widowed* | 3,812 | 25.6 |
| Social class |  |  |
| Managerial | 5,882 | 39.4 |
| Intermediate | 3,544 | 23.8 |
| Routine manual* | 5,488 | 36.8 |
| Wealth |  |  |
| Poorest* | 4,198 | 28.2 |
| Middle | 5,139 | 34.51 |
| Wealthiest | 5,577 | 37.4 |
| Education |  |  |
| Less than high school* | 4,619 | 30.9 |
| High school | 5,737 | 38.5 |
| Some college | 4,558 | 30.6 |

To gauge cohort effect initially, we computed proportions for all cohorts of people who were physically impaired based on thresholds recently established for the British population, i.e. 1.5 standard deviation below the mean (sex and yearly-age standardised) [4]. The cross-cohort proportions in Table 2 present, with a slight tapering off, a gradient of improvement as shown in reduction in impairment across cohort. Nearly 4.5% of Pre-Depression cohort members were impaired compared to only 2.4% of the Post-War cohort members.

**Table 2.** Proportion of physically impaired older people in each cohort based on British population reference values. Source: ELSA 2004-2013.

| Cohort | Impaired |  |
| --- | --- | --- |
|  | Yes | No |
| Pre-Depression | 4.45 | 95.55 |
| Depression cohort | 3.17 | 96.83 |
| War cohort | 2.19 | 97.81 |
| Post-War | 2.36 | 97.64 |
| Total | 2.77 | 97.23 |

To gain better insights we included other covariates in models of physical function. Model fit statistics, presented in Table 3, suggest that, as with Danish data [29], non-linear age trajectories fit the data best (smallest BIC and largest R^2^). Subsequent presentation relies on these four models (two outcomes for both sexes) with age and squared age terms.

**Table 3.** Model fit for gait speed and grip strength. Source: ELSA 2004–2013.

|  |  | Age |  | Age<sup>2</sup> |  |
| --- | --- | --- | --- | --- | --- |
|  |  | BIC | R<sup>2</sup> | BIC | R<sup>2</sup> |
| Women | Gait speed | 2121.794 | 0.434 | 1707.843 | 0.705 |
| Men |  | 956.636 | 0.720 | 598.624 | 0.735 |
| Women | Grip strength | 55612.431 | 0.127 | 55612.656 | 0.569 |
| Men |  | 47187.521 | 0.788 | 47185.125 | 0.789 |

The coefficients of the best models for gait speed are given in Table 4 and for grip strength in Table 5. Table 4 showed that there is no difference in gait speed at normal pace among those who were born during different times in the last century. Compared to those born before 1930 (the reference), men and women of the three subsequent cohorts of Depression era, War and Post-War cohorts walked no faster. Both sexes however showed similar reduction in gait speed with high levels of C-reactive protein: women by 0.023 m/s (95% confidence interval, CI: 0.037 – 0.090m/s) and men by 0.027 (CI: 0.011 – 0.043 m/s).

**Table 4.** Age trajectories of gait speed among Britons aged 50 years and older. Source: ELSA 2004 – 2013

| Gait speed | Women |  |  | Men |  |  |
| --- | --- | --- | --- | --- | --- | --- |
|  | coef | s.e. | p | coef | s.e. | p |
| Constant | 7.301 | 0.203 | < 0.001 | 7.241 | 0.216 | < 0.001 |
| Age | -0.160 | 0.006 | < 0.001 | -0.158 | 0.007 | < 0.001 |
| Age2 | 0.098 | 0.005 | < 0.001 | 0.097 | 0.005 | < 0.001 |
| Cohort: Pre-Depression as reference |  |  |  |  |  |  |
| Depression era | 0.021 | 0.021 | 0.31 | 0.016 | 0.023 | 0.47 |
| War | -0.042 | 0.027 | 0.11 | -0.047 | 0.029 | 0.10 |
| Post-War | 0.036 | 0.032 | 0.26 | 0.010 | 0.035 | 0.77 |
| Height | -0.000 | 0.000 | 0.001 | -0.000 | 0.000 | 0.008 |
| Wealth: poorest third as ref. |  |  |  |  |  |  |
| Middle | 0.013 | 0.008 | 0.098 | 0.003 | 0.008 | 0.70 |
| Wealthiest | 0.042 | 0.008 | < 0.001 | 0.030 | 0.009 | < 0.001 |
| Education: primary school as ref. |  |  |  |  |  |  |
| High school | 0.031 | 0.008 | < 0.001 | 0.039 | 0.009 | < 0.001 |
| College | 0.039 | 0.011 | < 0.001 | 0.056 | 0.010 | < 0.001 |
| Occupation: routine manual as ref. |  |  |  |  |  |  |
| Managerial | 0.028 | 0.010 | 0.003 | 0.024 | 0.008 | 0.003 |
| Intermed | 0.016 | 0.008 | 0.047 | 0.015 | 0.012 | 0.22 |
| Marital status: separated/widowed as ref. |  |  |  |  |  |  |
| Single | 0.006 | 0.016 | 0.71 | 0.002 | 0.015 | 0.90 |
| Married | -0.000 | 0.007 | 1.00 | 0.012 | 0.009 | 0.22 |
| High CRP | -0.023 | 0.007 | 0.002 | -0.027 | 0.008 | < 0.001 |
| High fibrinogen | -0.008 | 0.007 | 0.26 | -0.005 | 0.008 | 0.56 |
| $\sigma_{\text{int}}^2$ | 0.025 | 0.028 | < 0.001 | -0.469 | 0.033 | < 0.001 |
| $\sigma_{\text{age}}^2$ | 0.000 | 0.000 | 1.00 | 0.000 | 0.000 | 1.00 |
| $\sigma_{\text{int,age}}$ | 0.001 | 0.000 | < 0.001 | 0.009 | 0.001 | < 0.001 |

**Table 5.**
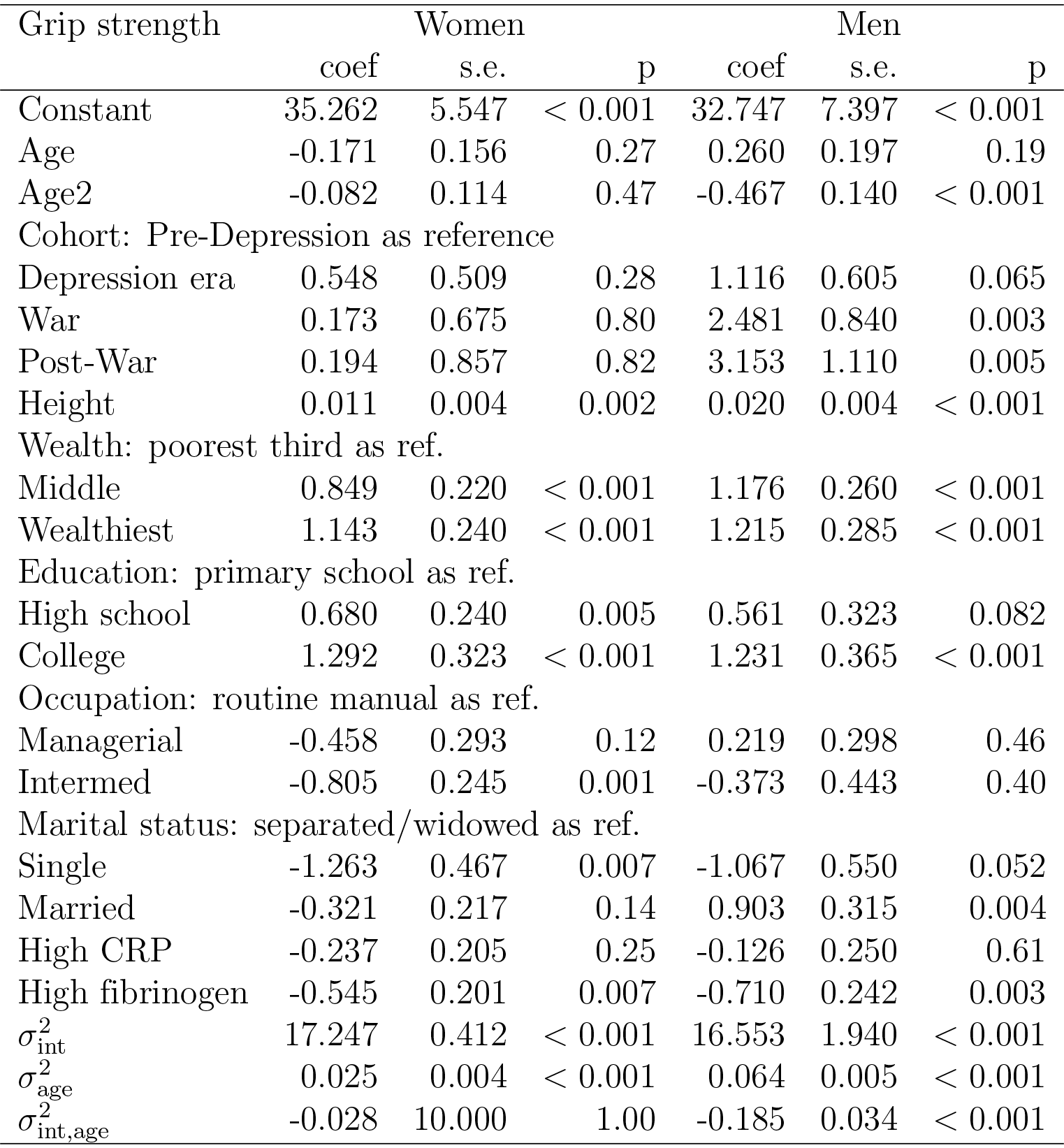
Age trajectories of grip strength among Britons aged 50 years and older. Source: ELSA 2004 – 2013

| Grip strength | Women |  |  | Men |  |  |
| --- | --- | --- | --- | --- | --- | --- |
|  | coef | s.e. | p | coef | s.e. | p |
| Constant | 35.262 | 5.547 | < 0.001 | 32.747 | 7.397 | < 0.001 |
| Age | -0.171 | 0.156 | 0.27 | 0.260 | 0.197 | 0.19 |
| Age2 | -0.082 | 0.114 | 0.47 | -0.467 | 0.140 | < 0.001 |
| Cohort: Pre-Depression as reference |  |  |  |  |  |  |
| Depression era | 0.548 | 0.509 | 0.28 | 1.116 | 0.605 | 0.065 |
| War | 0.173 | 0.675 | 0.80 | 2.481 | 0.840 | 0.003 |
| Post-War | 0.194 | 0.857 | 0.82 | 3.153 | 1.110 | 0.005 |
| Height | 0.011 | 0.004 | 0.002 | 0.020 | 0.004 | < 0.001 |
| Wealth: poorest third as ref. |  |  |  |  |  |  |
| Middle | 0.849 | 0.220 | < 0.001 | 1.176 | 0.260 | < 0.001 |
| Wealthiest | 1.143 | 0.240 | < 0.001 | 1.215 | 0.285 | < 0.001 |
| Education: primary school as ref. |  |  |  |  |  |  |
| High school | 0.680 | 0.240 | 0.005 | 0.561 | 0.323 | 0.082 |
| College | 1.292 | 0.323 | < 0.001 | 1.231 | 0.365 | < 0.001 |
| Occupation: routine manual as ref. |  |  |  |  |  |  |
| Managerial | -0.458 | 0.293 | 0.12 | 0.219 | 0.298 | 0.46 |
| Intermed | -0.805 | 0.245 | 0.001 | -0.373 | 0.443 | 0.40 |
| Marital status: separated/widowed as ref. |  |  |  |  |  |  |
| Single | -1.263 | 0.467 | 0.007 | -1.067 | 0.550 | 0.052 |
| Married | -0.321 | 0.217 | 0.14 | 0.903 | 0.315 | 0.004 |
| High CRP | -0.237 | 0.205 | 0.25 | -0.126 | 0.250 | 0.61 |
| High fibrinogen | -0.545 | 0.201 | 0.007 | -0.710 | 0.242 | 0.003 |
| $\sigma_{\text{int}}^2$ | 17.247 | 0.412 | < 0.001 | 16.553 | 1.940 | < 0.001 |
| $\sigma_{\text{age}}^2$ | 0.025 | 0.004 | < 0.001 | 0.064 | 0.005 | < 0.001 |
| $\sigma_{\text{int,age}}^2$ | -0.028 | 10.000 | 1.00 | -0.185 | 0.034 | < 0.001 |

Compared to the estimates for gait speed, those for maximum grip strength showed similarities and differences (Table 5). There is a discernible and different pattern of cohort effect. Men of the three subsequent cohorts displayed a step by step increase in maximum grip strength by about one kg per cohort, amounting to a cohort gradient or secular improvement. For the two most recent cohorts, this improvement is also highly statistically significant. There is also some similarity with the results on gait speed (Table 4) in the inverse association between inflammation and grip strength. However, this time the significant marker is fibrinogen. High levels of fibrinogen are associated with more than half a kilogram reduction in grip strength (women: 545 gram, CI: 151 – 939 gram; men: 710 gram, CI: 236 – 1,184 gram).

Other findings can be briefly summarised. Socioeconomic positions throughout the life course as indicated by wealth, occupation and education showed largely significant associations with both measures of physical function. Wealth (wealthiest and middle compared to the poorest third), occupation (managerial and intermediate compared to routine manual occupation), and education (college and high school compared to up to primary school leavers) have positive associations with both gait speed and grip strength, and are mostly statistically significant. A minor exception is noted where among women, intermediate occupation has a significantly negative coefficient compared to routine manual occupation. This may be due to more use of physical exertion in the routine manual occupation.

Finally, to illustrate the contributions of all covariates to grip strength, we plot predicted values of grip strength in Figure 1. We refrained from presenting an analogous plot for gait speed since cohort indicators were not found significant; and from commenting on the shapes of the trajectories in Figure 1, relying on fit statistics in Table 3 to decide on the best model. In Figure 3, the four cohorts of men, marked with (M), are above the four cohorts of women. Moreover, the War cohort (M) and the Post-War cohort (M) can be seen to be slightly above the older two cohorts (the Depression era cohort (M) and the Pre-Depression era cohort (M)). The statistical significance of the higher values should be gathered not from this figure but from Table 5, which suggest that the two most recent cohorts of men attained significantly higher values. In contrast, in the women’s sample there was no discernible difference in the four plots (clustered at the lower part), consistent with the lack of statistical significance shown in Table 5.

**Figure 1.**
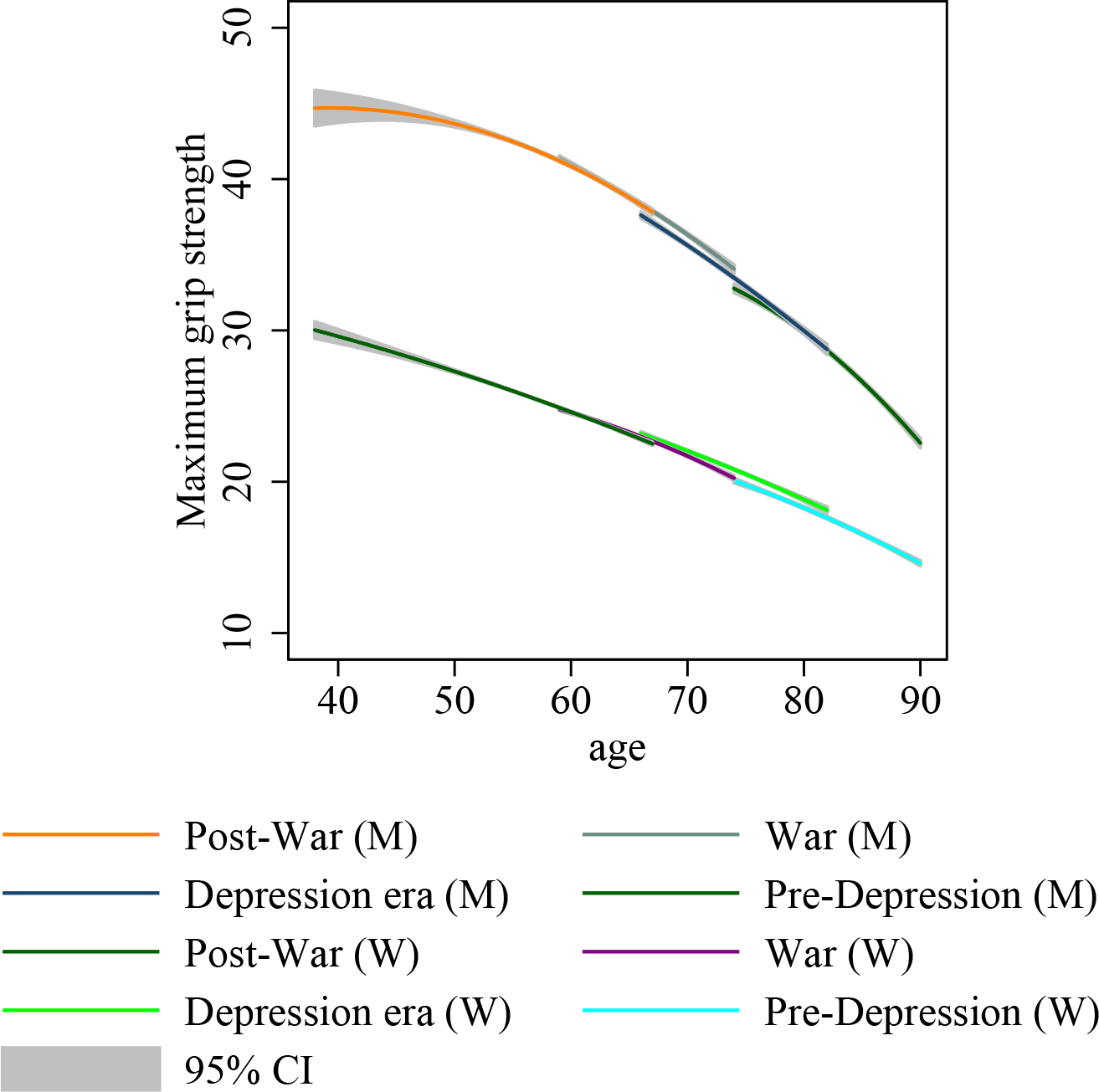
Predicted grip strength based on the best models. Source: ELSA 2004 – 2013.

## Discussion

The trend of physical disability in older people, with its cost implications, has been uncertain given the countervailing drivers of extending life expectancy and reduction in disability at a given age [10, 37]. Our analysis uncovered a secular improvement in physical function that is most pronounced among men born during and after the War. Unfortunately, no evidence of comparable gains accrued to women.

The data also revealed an intriguing pattern of improvement across physical function. The pattern is distinguished along normal capacity (gait speed at normal pace) versus maximum capacity (maximum grip strength). Among men, normal functional capacity did not show cross-cohort improvement at all but maximum capacity showed a secular improvement. The maximum force that muscles can physically muster when called forth has evidently increased across cohort considerably. This calls to mind that, in parallel, cognitive function in this sample has also been shown to improve across cohort [1]. This is the first evidence of a complex pattern of improvement in trajectories of physical function across cohort and sex.

The mechanism driving the cross-cohort improvement to health functioning has generally been ascribed to general improvement in public health infrastructure and education [5, 38, 39]. Improvements in public health infrastructure from the early part of the last century meant that children grew up with better conditions and reduced hazards and damage to health and child development. Improvements in education up to tertiary levels meant that adults became better equipped to make use of the new information that was abundantly created and increasingly available through the parallel progress in science and medical technology. Although such developments have not resulted in uniform and secular improvement in physical function, in maximum grip strength they have. Therein also lies a potential resolution to the uncertain trends in physical disability, i.e. different aspects of physical function give different pictures but maximum grip strength shows secular improvement.

A ground for optimism is thus available based on secular improvement in maximum capacity among men. But as noted in the WHO World Report on Ageing and Health [10] the daily functioning of an older person crucially depends on the surrounding health system environments and on access to such systems. Two older persons of the same cohort with similarly low level of maximum capacity may fare and function differently depending on their access to assistive technology to compensate for the perceived gap. Nevertheless, given the secular improvement in both cognitive and physical functions among men, the more recent cohorts of the older population hold a double potential for continuing contribution that may not have been fully appreciated.

Inflammation, on the other hand, is largely harmful across both measures of health function. Although different markers are found to be significant for different measures, inflammation is inversely related to maximum and normal functional capacity. Thus high fibrinogen associates with weaker grip while high CRP associates with slower gait. This gives some contrast to previous work on this sample. In a cross-section study of average grip strength, CRP has been found to be significant [40]. Our longitudinal study showed a similar sign but not significance. In comparison with a cross-sectional observation, longitudinal observations which were analysed with due control for attrition offer some advantage, particularly control for unobserved individual differences.

The inflammation effect echoes a finding based on this sample which showed inflammation to be harmful to cognitive function [15]. Evidently, inflammation also goes with reduced physical function, supporting the idea of inflammaging [41]. The mechanism for this revolves around the role of inflammatory cytokines in both muscle regeneration and muscle functioning. In normal activities of daily living which involve muscle exertion, some minute damage to muscle tissue may occur. In these circumstances, the pluripotent myosatellite cells respond by proliferating and differentiating to form muscle fibres and cover the damaged tissue. Circulating inflammatory cytokines such as tumour necrosis factor *α* (TNF*α*) have been shown to impair this process of regeneration in two ways: apoptosis of myosatellite cells and inhibition of the differentiation stage, leaving proliferated cells unable to differentiate and replace the damaged tissue. Beyond impairing the myogenesis process in common minute damage, inflammation also impairs functioning by reducing the power of the single permeable fibre. So in mice, TNF*α* rapidly reduces the force generating capacity or specific tension of muscle fibres independent of loss of muscle volume. In short, inflammation impairs muscle functioning in older people in at least three ways: it encourages myosatellite cell deaths, it interrupts the step of differentiation into myonuclei and muscle fibres; lastly, even if muscle fibres have been successfully regenerated, inflammation reduces the febrile tensile strength.

This study has a number of weaknesses. First, not all common measures of inflammatory cytokines were collected, especially TNF*α*. Addressing this should help in strengthening the mechanism by securing close comparison between population studies and *in vitro* studies. Since muscle strength is determined to a large extent by muscle volume [42], a better measure of muscle volume using dual energy x-ray absorptiometry can additionally strengthen the basis for the mechanism underlying the observed improvement. Lastly, the complex result on cohort improvement, depending on aspects of physical function and sex, may be highly specific to the British experience. A cross-country comparison is an obvious next step. This study nonetheless has some strengths. First, the sample is designed to represent the country and not only some clinical groups or regions, hence facilitating generalisation. Moreover, this is the first study, based on repeated measures of both physical function and inflammation, to draw trajectories of physical function and factors that shaped them as they unfold with age. Finally, this study also derived the trajectories while dealing with the attrition that is common but often ignored in longitudinal ageing studies.

In conclusion, pronouncements about trends in healthy physical ageing are marked with inconsistency [43] and some confusion [10]. Recent results on cognitive ageing in Britain are reinforced with these newly uncovered results: among men both cognitive and physical functions are secularly improving. Future responses to the challenge of an ageing population [9] by research and policy should carefully consider cohort composition to gain useful insights and craft efficient policy.

## Acknowledgments

This work was supported by grants from the Medical Research Council and Economic & Social Research Council (No. G1001375/1), the National Institute of Health Research (No. ES/L001772/1) and the European Union’s Horizon 2020 Research and Innovation Programme (No. 668648) to Gindo Tampubolon.

